# Surface association induces cytotoxic alkyl-quinolones in *Pseudomonas aeruginosa*

**DOI:** 10.1101/773291

**Authors:** Geoffrey D. Vrla, Mark Esposito, Chen Zhang, Yibin Kang, Mohammad R. Seyedsayamdost, Zemer Gitai

## Abstract

Surface attachment, an early step in the colonization of multiple host environments, activates the virulence of the human pathogen *P. aeruginosa*. However, the signaling pathways and downstream toxins specifically induced by surface association to stimulate *P. aeruginosa* virulence are not fully understood. Here, we demonstrate that alkyl-quinolone (AQ) secondary metabolites are rapidly induced upon surface association and represent a major class of surface-dependent cytotoxins. AQ cytotoxicity is direct and independent of other AQ functions like quorum sensing or PQS-specific activities like iron sequestration. Furthermore, the regulation of AQ production can explain the surface-dependent virulence regulation of the quorum sensing receptor, LasR, and the pilin-associated candidate mechanosensor, PilY1. PilY1 regulates surface-induced AQ production by repressing the AlgR-AlgZ two-component system. AQs also contribute to the known cytotoxicity of secreted outer membrane vesicles. These findings collectively explain previously mysterious aspects of virulence regulation and provide new avenues for the development of anti-infectives.

## Introduction

The opportunistic human pathogen *P. aeruginosa* infects a wide range of hosts such as mammals, plants, insects, and fungi (***Rahme et al., 1995***), and is a major contributor to the morbidity of cystic fibrosis patients (***Nixon et al., 2001***) and hospital-acquired infections (***Richards et al., 1999***). *P. aeruginosa* uses a large set of secreted proteins and secondary metabolites to carry out the multiple requirements necessary for a successful infection, including host colonization, immune evasion, nutrient acquisition, and host cell killing (cytotoxicity) (***Valentini et al., 2018***). Given the multiple activities involved in pathogenesis, we recently developed a quantitative imaging-based host cell killing assay to specifically study the factors acutely required for killing host cells during short timescales (***Siryaporn et al., 2014***). This assay revealed that cytotoxicity is activated by attachment of *P. aeruginosa* to a solid surface (***Siryaporn et al., 2014***). This surface-induced cytotoxicity does not require the Type-IV Pilus (TFP), TFP-associated signaling complexes (PilA-Chp-Vfr/cAMP), or Type III Secretion Systems (T3SS), but does require two regulatory proteins, LasR and PilY1 (***Siryaporn et al., 2014***). Since well-characterized cytotoxins such as T3SS and Vfr targets are not necessary for surface-induced host-cell killing in this assay, we sought to address the outstanding questions of which specific toxins mediate host cell killing in response to surface attachment and how these toxins are regulated by LasR and PilY1.

LasR is an important component of the complex network of *P. aeruginosa* quorum sensing (***Lee and Zhang, 2015***). Quorum sensing (QS) is the process by which bacteria synthesize and secrete autoinducer signaling molecules that accumulate and activate their receptors as a function of bacterial cell density. There are at least three QS systems that have been previously implicated in regulating *P. aeruginosa* virulence: the *las, rhl*, and *pqs* QS systems (***Lee and Zhang, 2015***). These systems form a complex and interconnected network with extensive regulatory cross-talk (***Maura et al., 2016***). For example, LasR transcriptionally upregulates the autoinducer synthase enzymes of the *rhl* and *pqs* systems (***Xiao et al., 2006b***; ***Farrow et al., 2008***), which in turn activate numerous downstream factors (***Farrow et al., 2008***). As a result, identifying the specific contribution of LasR dependent QS to the phenomenon of surface-induced virulence has been challenging.

Besides LasR, the other factor known to be required for surface-induced virulence is PilY1 (***Sirya-porn et al., 2014***), a minor pilin-associated protein with a putative mechanosensory domain (***Bohn et al., 2009***). PilY1 promotes several surface-dependent behaviors including virulence induction (***Siryaporn et al., 2014***), twitching motility (***Bohn et al., 2009***), and biofilm formation (***Kuchma et al., 2015***; ***Luo et al., 2015***). Like QS, there are multiple signaling pathways that have been proposed to function downstream of PilY1 (***Luo et al., 2015***). However, disrupting the key effectors of the two best-characterized pathways downstream of PilY1, c-di-GMP or cAMP production, does not influence surface-induced virulence (***Siryaporn et al., 2014***).

Understanding the signaling pathways that LasR and PilY1 use to trigger surface-induced virulence has been particularly challenging because the relevant output of these pathways, such as the cytotoxin(s) that *P. aeruginosa* secretes to kill host cells when surface-associated, has remained unknown. *P. aeruginosa* possesses numerous candidate toxins that could mediate surface-induced virulence, including the type III secretion system (T3SS) and numerous other secreted proteins and secondary metabolites (***Valentini et al., 2018***). Many of these candidates were previously found not to be required for surface-induced virulence (***Siryaporn et al., 2014***), which could reflect functional redundancy or the existence of a previously-overlooked cytotoxin.

Here we identify the pathways that activate surface-induced virulence by first showing that a single family of cytotoxins, the alkyl-quinolones (AQs), are both necessary and sufficient to explain the surface-induced killing of *Dictyostelium discoideum* by *P. aeruginosa*. We demonstrate that surface association triggers increased AQ secretion, which requires both LasR and PilY1. We show that these findings are also relevant to mammalian host cells and demonstrate that AQs are a major cytotoxic component of secreted outer membrane vesicles. Together our data support the conclusion that surface-induced virulence results from induction of AQs, which act as toxins that directly kill host cells.

## Results

### Alkyl-quinolones are necessary and sufficient for surface-induced virulence of *P. aeruginosa* towards *D. discoideum*

Surface attachment strongly stimulates the ability of *P. aeruginosa* PA14 to kill *D. discoideum* amoebae (***Siryaporn et al., 2014***). Time-lapse imaging of *D. discoideum* infected with planktonic *P. aeruginosa* demonstrates that *D. discoideum* completely clears the bacterial population through phagocytosis (Figure 1-S1). Similar treatment of *D. discoideum* with *P. aeruginosa* that had been previously attached to a glass surface results in reduced *D. discoideum* motility and eventually leads to cell lysis (Figure 1-S2). To identify the factors required for the increased virulence of surface-associated *P. aeruginosa* we screened a number of mutants in secreted effectors known to promote pathogenesis for loss of virulence following surface attachment (Figure 1-S3). Specifically, we grew each mutant to the same density, allowed it to associate with a glass surface for 1 hour, added *D. discoideum* host cells, and monitored host cell death by fluorescence microscopy using the live-cell-impermeant dye, calcein-AM. Loss of many candidate *P. aeruginosa* cytotoxins or virulence regulators did not reduce surface-induced killing of *D. discoideum*, including phenazines, rhamnolipids, and hydrogen cyanide (Figure 1-S3). In contrast, *pqsA* was absolutely required for surface-induced virulence (Figure 1A). PqsA is an enzyme required for the biosynthesis of alkyl-quinolones (AQs) such as PQS, HHQ, and HQNO (***Coleman et al., 2008***), suggesting that AQs play a key role in surface-induced virulence.

The alkyl-quinolone (AQ) family of small molecules in *P. aeruginosa* performs a diverse set of virulence-related functions including quorum-sensing signaling (***Rampioni et al., 2016***), iron acquisition (***Diggle et al., 2007***), immune suppression (***Kim et al., 2010***), and anti-bacterial activities (***Hazan et al., 2016***). One AQ species, PQS, has been suggested to possess cytotoxic activity towards some mammalian cell types (***Abdalla et al., 2017***) and HQNO can disrupt electron transport in vitro (***Reil et al., 1997***). To identify the parts of the AQ pathway responsible for surface-induced virulence of *P. aeruginosa* towards *D. discoideum*, we assayed mutants deficient in the four keys steps of the AQ pathway (Figure 1B): 1) converting the anthranilate precursor to HHQ (mediated by PqsABCD), 2) converting HHQ into PQS and HQNO (by PqsH and PqsL, respectively), 3) feedback regulation onto *pqsABCDE* expression (by PQS and HHQ activating the transcriptional regulator PqsR, also known as MvfR), and 4) stimulating RhlR-dependent QS (by PqsE). Neither the production of HQNO (Δ*pqsL*) nor the activation of rhlR-dependent targets (Δ*pqsE*) was required for the killing of *D. discoideum* by *P. aeruginosa* (Figure 1C). However, *pqsA, pqsH* and *pqsR* mutants showed reduced ability to kill *D. discoideum* (Figure 1C).

**Figure 1.**
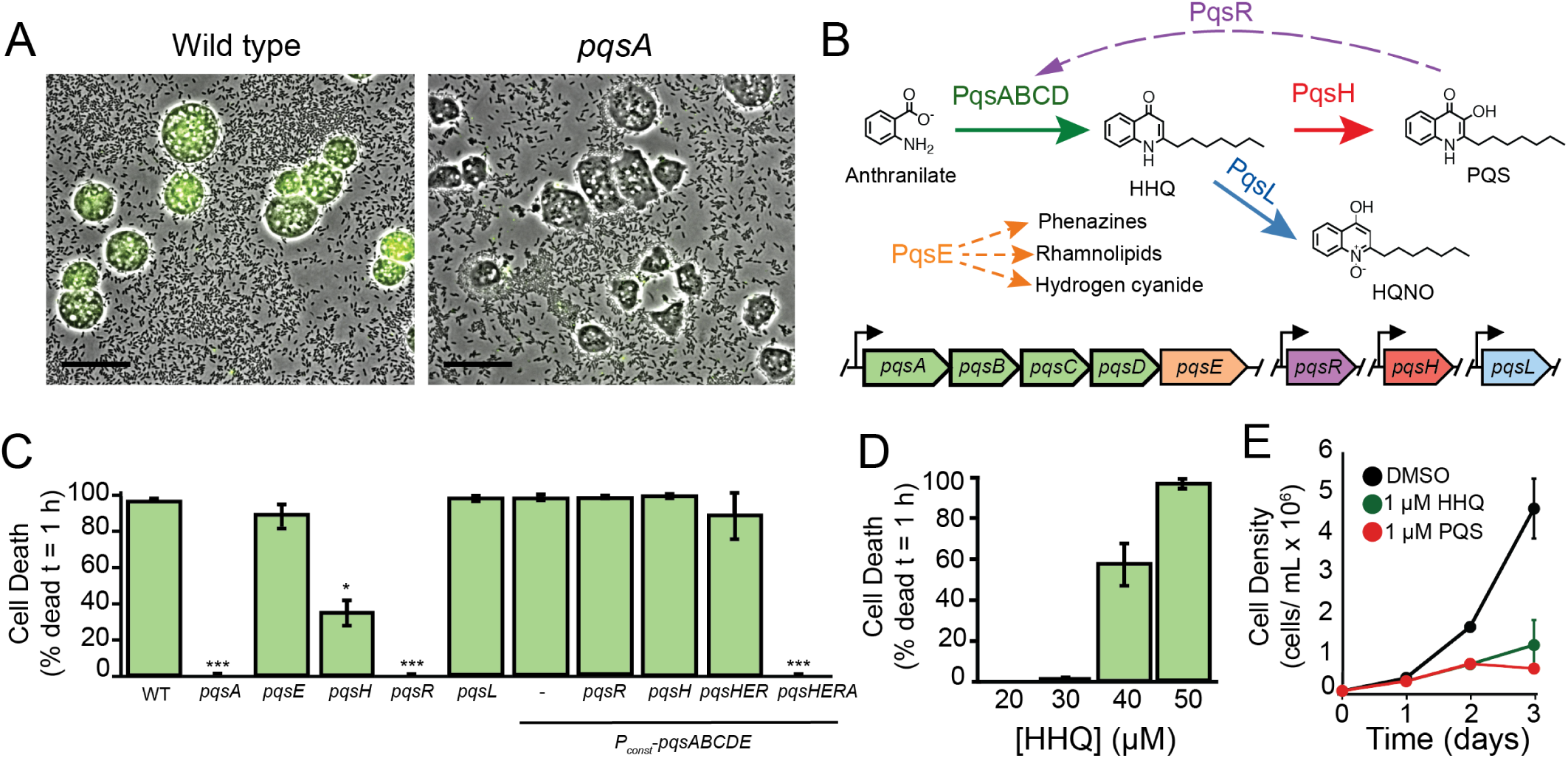
Alkyl-quinolone production is necessary and sufficient for surface-induced killing of *D. discoideum* by *P. aeruginosa*. (A) *D. discoideum* infected with surface-attached wild type and Δ*pqsA P. aeruginosa* after 1 h co-culture (scale bars = 30 *μ*m). Fluorescent calcein-AM staining indicates cell death. (B) Schematic of the PQS pathway depicting the functions of relevant genes. Solid and dotted arrows represent biosynthetic reactions and gene regulation, respectively. (C) Quantification of *D. discoideum* killing by PQS pathway mutants. Expression of *pqsABCDE* is driven by the endogenous *pqsA* promoter or a strong constitutive promoter inserted upstream of the *pqsA* gene (*P*_*const*_ − *pqsABCDE*) (D) Cytotoxicity of purified HHQ to *D. discoideum* under conditions of the cell death assay in (C). (E) Axenic growth of *D. discoideum* in the presence of purified HHQ or PQS. Values are mean ± SD of five biological replicates (n = 5). Values in (C) and (D) are mean ± SEM of three biological replicates (n = 3). For statistical analysis in (C), mutants were compared to wild type (Student’s t-test, two-tailed, * = p<0.05, ***p<0.001). Approximately 200-300 cells were analyzed for each measurement in (C) and (D). **Figure 1–Figure supplement 1.** Time-lapse imaging of *D. discoideum* infected with planktonic *P. aeruginosa* **Figure 1–Figure supplement 2.** Time-lapse imaging of *D. discoideum* infected with surface-attached *P. aeruginosa* **Figure 1–Figure supplement 3.** Virulence of various *P. aeruginosa* mutants towards *D. discoideum*. **Figure 1–Figure supplement 4.** Virulence of *pqsA* and *pqsH* mutants supplemented with HHQ. **Figure 1–Figure supplement 5.** Quantification of AQ production in *P*_*const*_ − *pqsABCDE* strains by LC/MS

The reduced virulence of Δ*pqsH* suggested that PQS contributes to surface-induced virulence, which could be due to its role in iron acquisition, its QS ability to activate PqsR, or its activity as a cytotoxin. To address the potential role of iron binding, we relied on the fact PQS binds iron while its HHQ precursor does not (***Diggle et al., 2007***). Deleting *pqsH*, the enzyme that converts HHQ to PQS, results in reduced levels of all AQs due to the absence of PqsR-mediated feedback induction. We thus transiently supplemented *pqsH* and *pqsA* mutants with HHQ for 1 hour at concentrations sufficient to induce PqsR (50 *μ*M). We then washed away the HHQ before exposing these bacteria to *D. discoideum*, such that during the virulence assay the *pqsA* mutants would have no HHQ or PQS while *pqsH* mutants would have HHQ but no PQS. We found that transient HHQ addition was sufficient to restore WT-levels of virulence to *pqsH* mutants but not to *pqsA* mutants (Figure 1-S4), indicating that HHQ is required for surface-induced virulence but PQS is not.

Having eliminated PQS-specific roles of AQs such as iron sequestration, we next tested if surface-induced virulence requires PqsR activation to achieve high levels of the AQs themselves or if additional PqsR targets (***Maura et al., 2016***) might be required. Consequently, we replaced the endogenous *pqsA* promoter with a strong constitutive promoter (*P*_*OXB20*_), which we refer to as (*P*_*const*_). This construct was sufficient to fully restore the virulence of *pqsR* and *pqsH* mutants to WT levels (Figure 1C). These results suggest that the requirement of PQS production for full virulence is due to the ability of PQS to promote high levels of *pqsABCDE* by activating the PqsR-dependent positive feedback loop. To further eliminate the possibility of PqsE-mediated QS, we simultaneously deleted *pqsE, pqsH*, and *pqsR* in the (*P*_*const*_ − *pqsABCDE*) background and showed that these bacteria retained PqsA-dependent virulence (Figure 1C). To confirm that each of the strains analyzed retained the expected ability to produce HHQ and/or PQS, we used LC/MS to directly quantify these molecules in the bacterial supernatants and found the expected results in all cases (Figure 1-S5).

The ability of high expression of *pqsABCDE* to induce virulence in the absence of *pqsH* suggested that surface-induced virulence is mediated either by HHQ itself, or by another previously uncharacterized product of *pqsABCDE*. To determine if HHQ is not just required, but also sufficient for killing *D. discoideum* we obtained commercially-purified HHQ (Sigma Aldrich, St. Louis, MO) and determined its lethal dose under the conditions used in the surface-induced virulence assay (Figure 1D). Purified HHQ induced rapid cell death in *D. discoideum* at concentrations above 30 *μ*M (Figure 1D), and inhibited growth of *D. discoideum* at concentrations as low as 1 *μ*M when added to axenic *D. discoideum* cultures (Figure 1E). While PQS was not necessary for virulence, PQS also acted as a direct cytotoxin as purified PQS killed *D. discoideum* (Figure 1E). Together, our results demonstrate that HHQ and PQS are cytotoxic towards *D. discoideum*, that AQ production is both necessary and sufficient for surface-induced virulence of *P. aeruginosa*, and that AQs other than PQS can mediate this toxicity.

### AQ regulation can explain the effects of known surface-induced virulence regulators

We previously showed that LasR and PilY1 are required for surface-induced virulence (***Siryaporn et al., 2014***). To understand whether the virulence defects of these mutants are due to loss of AQ production, we determined if they can be rescued by replacing the endogenous *pqsA* promoter with the *P*_*OXB20*_ strong constitutive promoter (*P*_*const*_). This *pqsABCDE* overexpression restored full virulence to surface-associated *lasR* and *pilY1* deletion mutants (Figures 2A and 2B). Furthermore, constitutive *pqsABCDE* expression was sufficient to induce detectable virulence in planktonic bacteria (Figures 2A and 2B). While the virulence achieved by expression of *pqsABCDE* in planktonic cells did not reach the same levels as the surface-attached bacteria, the conversion of avirulent cells to a virulence state with only the expression of a single operon is notable.

**Figure 2.**
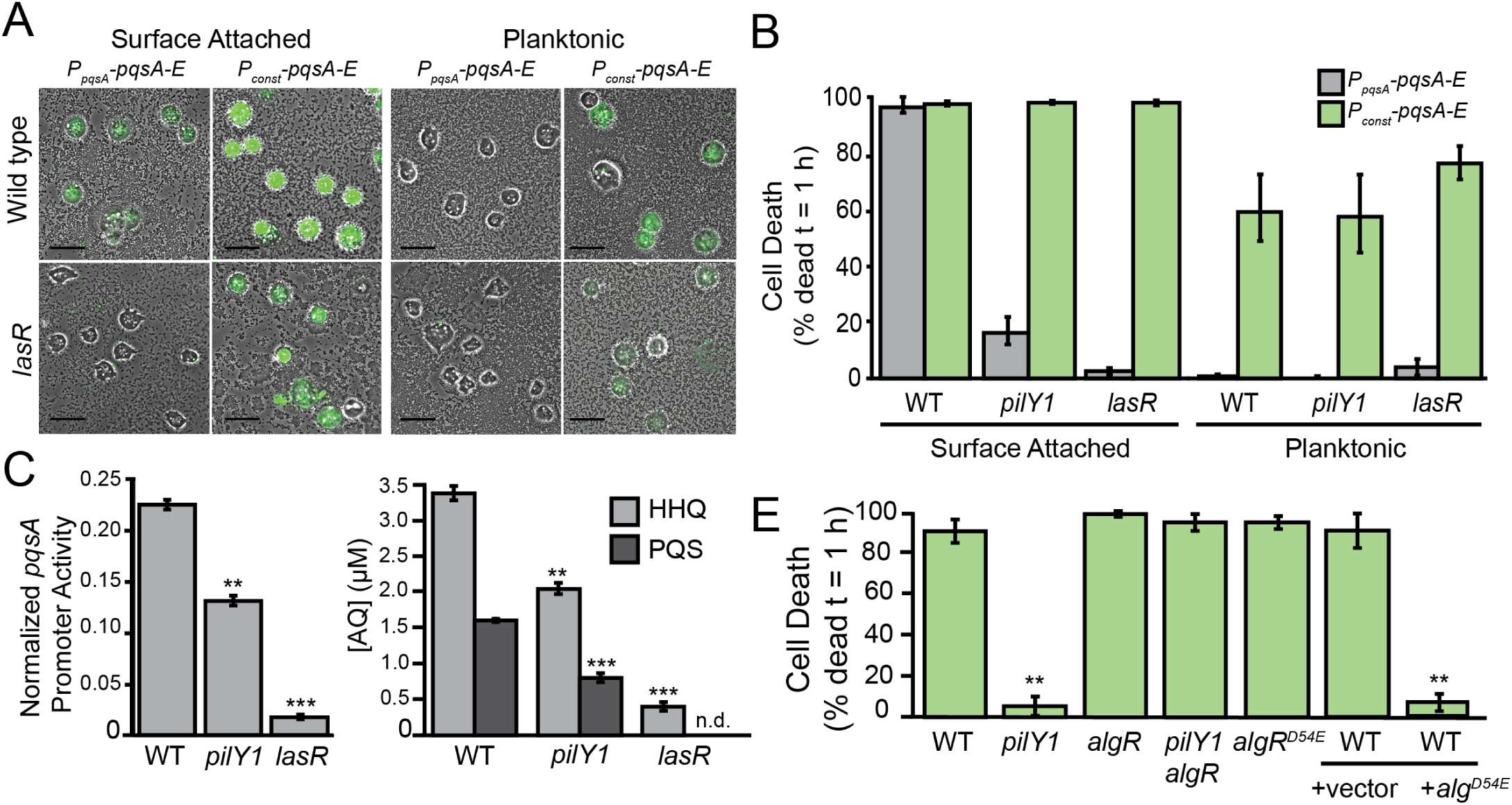
PilY1 and LasR promote surface-induced virulence through *pqsABCDE* expression. (A) Representative images of *D. discoideum* infected with surface-attached and planktonic *P. aeruginosa* expressing *pqsABCDE* genes under control of the endogenous *pqsA* promoter (*P*_*pqsA*_ − *pqsABCDE*) or a strong constitutive promoter (*P*_*const*_ − *pqsABCDE*) after 1 hour of co-culture (scales bars = 30 *μ*m). Fluorescent calcein-AM staining indicates cell death. (B) Quantification *D. discoideum* killing by surface-attached and planktonic wild type, Δ*pilY1* and Δ*lasR P. aeruginosa* after 1 h co-culture. (C) Mean fluorescence intensity per cell (>500 cells) of surface-attached *P. aeruginosa* expressing a (*P*_*pqsA*_ − *mCherry*) promoter fusion. Values are mean ± SEM of three biological replicates (n = 3). (D) LC/MS-based quantification of HHQ and PQS in extracts of wild type, Δ*pilY1*, and Δ*lasR* liquid cultures grown to OD_600*nm*_ = 1.5. Values are mean ± SEM of three biological replicates (n = 3), and concentrations were calculated using a standard curve constructed from purified AQ standards (n.d.= not detected). (E) Quantification *D. discoideum* killing by mutants in the AlgR-PilY1 pathway after 1 h co-culture. Values in (B) and (E) are mean ± SEM of three biological replicates (n = 3). Statistical analysis of mutants in (C) were performed against wild type (Student’s t-test, two-tailed, * = p<0.05, ** p <0.01, ***p<0.001). Approximately 150-300 cells were analyzed for each measurement in (B) and (E).

We next sought to determine if *pqsABCDE* expression is altered in *lasR* and *pilY1* mutants using a fluorescent reporter fusion to the *pqsABCDE* promoter. We fused a 500-bp fragment upstream of the *pqsA* gene to a promoterless *mCherry* gene, and integrated this construct at a neutral chromosomal locus. Wild type, Δ*lasR*, and Δ*pilY1* bacteria expressing this reporter were grown to mid-exponential phase (OD_600*nm*_ = 0.6), and OD-matched cultures were allowed to attach to a surface for 1 hour. We measured the mean fluorescence intensity of individual surface-attached bacteria and normalized them to a constitutively expressed fluorescent reporter. The *pqsABCDE* promoter activity of both Δ*lasR* and Δ*pilY1* was significantly lower than that of wild type (Figure 2C). To determine if the decreased promoter activity impacts AQ production, we measured HHQ and PQS levels in ethyl acetate extracts of wild type, Δ*lasR*, and Δ*pilY1* cultures grown to early stationary phase using LC/MS. This revealed that both HHQ and PQS production is decreased in both Δ*lasR* and Δ*pilY1* compared to wild type (Figure 2D). Together these results suggest that the effects of known virulence regulators can be explained by their effects on *pqsABCDE* operon expression.

LasR has previously been show to induce the PqsR *pqsABCDE* regulator (***Xiao et al., 2006b***), but the connection between PilY1 and *pqsABCDE* expression has not been previously reported. Since the well-characterized c-di-GMP and cAMP pathways do not appear responsible for PilY1-mediated virulence induction (***Siryaporn et al., 2014***), we compared the previously-reported surface-dependent transcriptional changes in WT and *pilY1* mutants (***Siryaporn et al., 2014***). We found that both the AlgR-AlgZ TCS and its associated regulon are repressed in a PilY1-dependent-manner (***Siryaporn et al., 2014***; ***Belete et al., 2008***). To test if AlgR can regulate virulence we generated a phosphomimetic mutation in *algR* (*algRD54E*) and overexpressed it from a multi-copy plasmid. AlgRD54E overexpression strongly inhibited virulence (Figure 2E). To determine if AlgR functions downstream of PilY1 we turned to epistasis analysis and generated a *pilY1 algR* double mutant, which restored virulence to the avirulent *pilY1* single mutant (Figure 2E). Thus, *algR* is epistatic to *pilY1* and likely functions downstream of it. To determine if PilY1 acts on the levels or phosphorylation state of AlgR, we replaced the chromosomal copy of *algR* with a phosphomimetic *algRD54E* allele. Unlike in the case of overexpressing *algRD54E* from a plasmid, chromosomal expression of *algRD54E* did not affect virulence (Figure 2E), which suggests that PilY1 functions by transcriptionally repressing the levels of *algR*.

### Virulent surface-attached cells secrete more AQs than avirulent planktonic cells

AQs have not been previously described to be surface-regulated, but our findings above suggested that surface attachment might stimulate AQ production. AQ quantification under the conditions of our surface-induced virulence assay is challenging using traditional MS-based techniques. Consequently, we developed a fluorescence-based AQ biosensor that can be used to measure PQS and HHQ levels under the exact conditions of the surface-induced *D. discoideum* cell death assay. Specifically, we engineered a reporter strain with three features: 1) it is unable to synthesize AQ’s itself (due to deletion of *pqsA*), 2) it linearly responds to PqsR activation without quorum-sensing feedback (due to replacement of the *pqsR* promoter with a constitutive *P*_*tac*_ promoter), and 3) it has a plasmid containing both a fluorescent YFP reporter for PqsR activation by AQs (*P*_*pqsA*_ − *YFP*) and a constitutive mKate reporter (*P*_*rpoD*_ − *mKate*) to normalize for plasmid copy number.

We validated our AQ biosensor by analyzing its response to purified AQ standards (Sigma-Aldrich, St. Louis, MO) in a 96-well format and by examining it in mutants with known effects on AQ production. The AQ biosensor responded linearly to PQS and HHQ across the range of AQ levels previously reported (***Xiao et al., 2006a***) for *P. aeruginosa* cultures (Figure 3A). Consistent with the known binding affinities of PqsR (***Xiao et al., 2006a***) the sensor responded more strongly to PQS than HHQ and did not respond to HQNO (Figure 3A and 3B). To use the biosensor to monitor AQ production by *P. aeruginosa* we doped the AQ reporter 1:100 into surface-attached wild type, Δ*pilY1*, Δ*algR*, and Δ*pilY1* Δ*algR*. We note that the AQ biosensor is itself avirulent such that doping it at low levels (1:100) enabled us to quantify AQ production without disrupting the assay. Consistent with the epistasis results obtained with respect to virulence (Figure 2E), the AQ reporter showed reduced AQ levels upon deletion of *pilY1*, and this decrease was suppressed in a *pilY1 algR* double mutant (Figure 3-S1).

**Figure 3.**
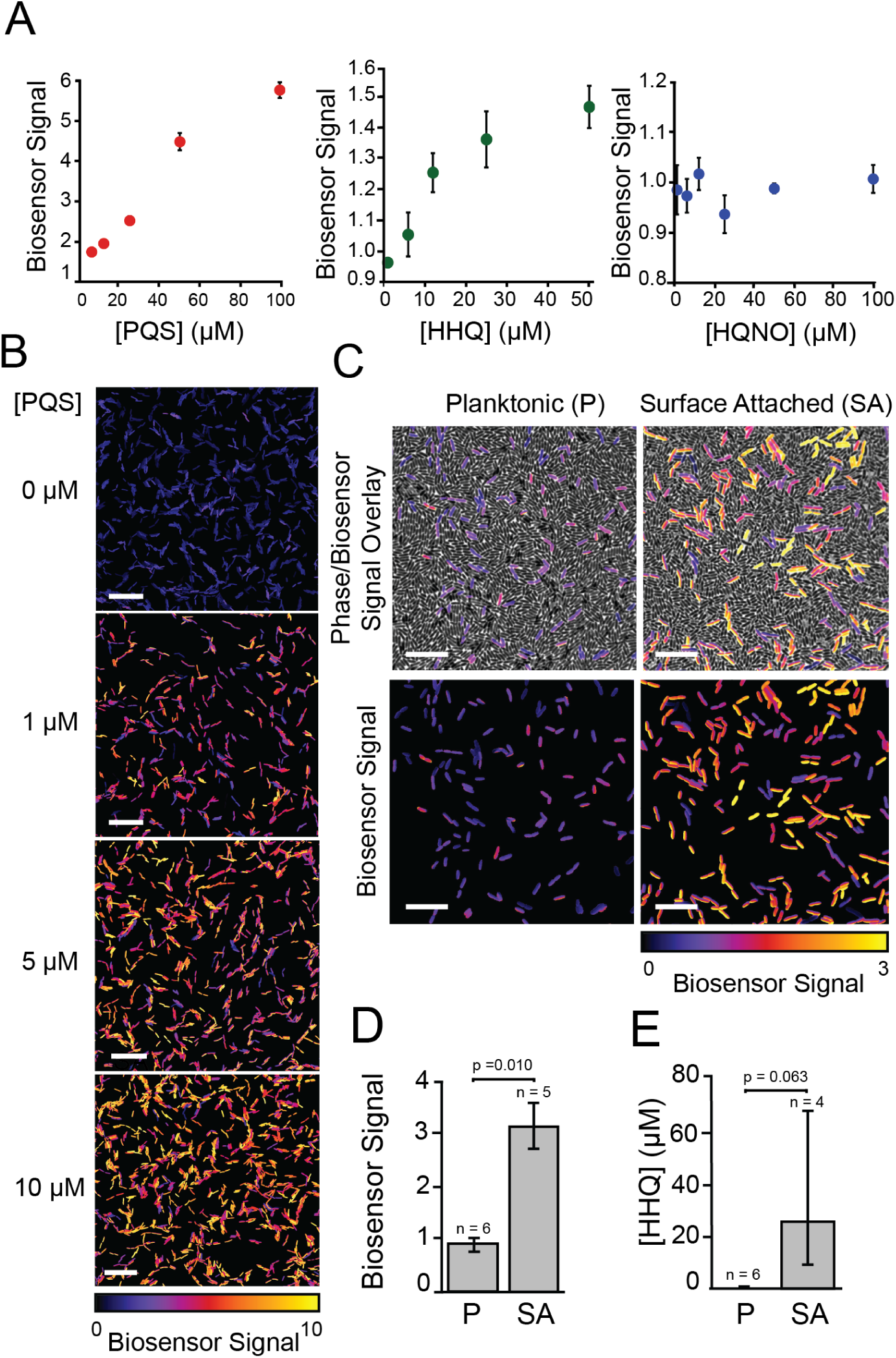
Biosensor-based detection of alkyl-quinolones in planktonic and surface-attached *P. aeruginosa* populations. (A) Response of AQ biosensor to increasing concentrations of purified PQS, HHQ, and HQNO in a 96-well microplate-based assay. Biosensor signal was calculated by normalizing YFP/mKate fluorescence by a DMSO control. Values are mean ± SD of six technical replicates (n = 6). (B) Representative images of biosensor signal of single cells in response to addition of purified PQS added to 1 % agar pads used for imaging (scale bars = 10 *μ*m). (C) Representative images of biosensor doped 1:100 into planktonic or surface-attached Δ*pqsH P. aeruginosa* grown under conditions of the *D. discoideum* cell death assay (scale bars = 5 *μ*m). Top images show an overlay of phase images and biosensor signal and bottom images show only the biosensor signal. (D) Quantification of biosensor signal in (C). Biosensor signal is the mean YFP/mKate fluorescence per cell (>500 cells) subtracted by the value from biosensor doped into surface-attached *pqsA* mutant, as describe in the material and methods. (E) HHQ concentration calculated from biosensor signal in (D) using an HHQ standard (Figure 3-S1). Values in (D) and (E) are mean ± SEM of indepdent biological replicates (n = 4-6). Statistical analysis in (D) is Student’s t-test (two-tailed, ** p <0.01). **Figure 3–Figure supplement 1.** Biosensor-based quantification of AQ production in *algR* mutants **Figure 3–Figure supplement 2.** HHQ standard curve for biosensor-based AQ quantification of surface-attached *P. aeruginosa*

Having validated our AQ biosensor, we used it to compare AQ levels between planktonic and surface-attached *P. aeruginosa* populations. Because the AQ biosensor responds to both HHQ and PQS, we focused on quantifying AQs from Δ*pqsH*, which makes HHQ but not PQS. We note that this strain is less virulent than wild type but retains 40% of its virulence and its virulence is still specifically induced by surface-association (Figure 1C). We doped the AQ biosensor (1:100) into cultures of Δ*pqsH* at the time of *D. discoideum* addition and observed a significant increase in biosensor signal as compared to planktonic Δ*pqsH* populations (Figures 3C and 3D). Conversion of biosensor signal to HHQ concentration using an HHQ standard curve (Figure 3-S2) indicated that surface-attached *pqsH* bacteria secrete at least 20-fold more HHQ than planktonic cells (Figure 3D).

### AQ cytotoxicity promotes surface-induced virulence towards mammalian cells and is important for outer membrane vesicle toxicity

To determine if our findings using *D. discoideum* host cells can be applied to mammalian hosts, we assayed the toxicity of purified HHQ, PQS, and HQNO towards adherent TIB-67 mouse monocytes and A549 human lung epithelial cells. Specifically, we added various concentrations of each purified AQ to the mammalian cells under standard culture conditions in a 96-well format and assessed viability after 48 hours using a water-soluble tetrazolium assay (Figure 4A). Both HHQ and PQS inhibited growth of TIB-67 in range of the concentrations observed in *P. aeruginosa* cultures by mass spectrometry, while HQNO did not (comparing Figure 4A to Figure 1-S3). PQS was significantly more toxic to TIB-67 than HHQ (Figure 4A). Both PQS and HHQ were also cytotoxic towards the A549 lung epithelial cell line, but this activity was lower than that against TIB-67 (Figure 4A).

**Figure 4.**
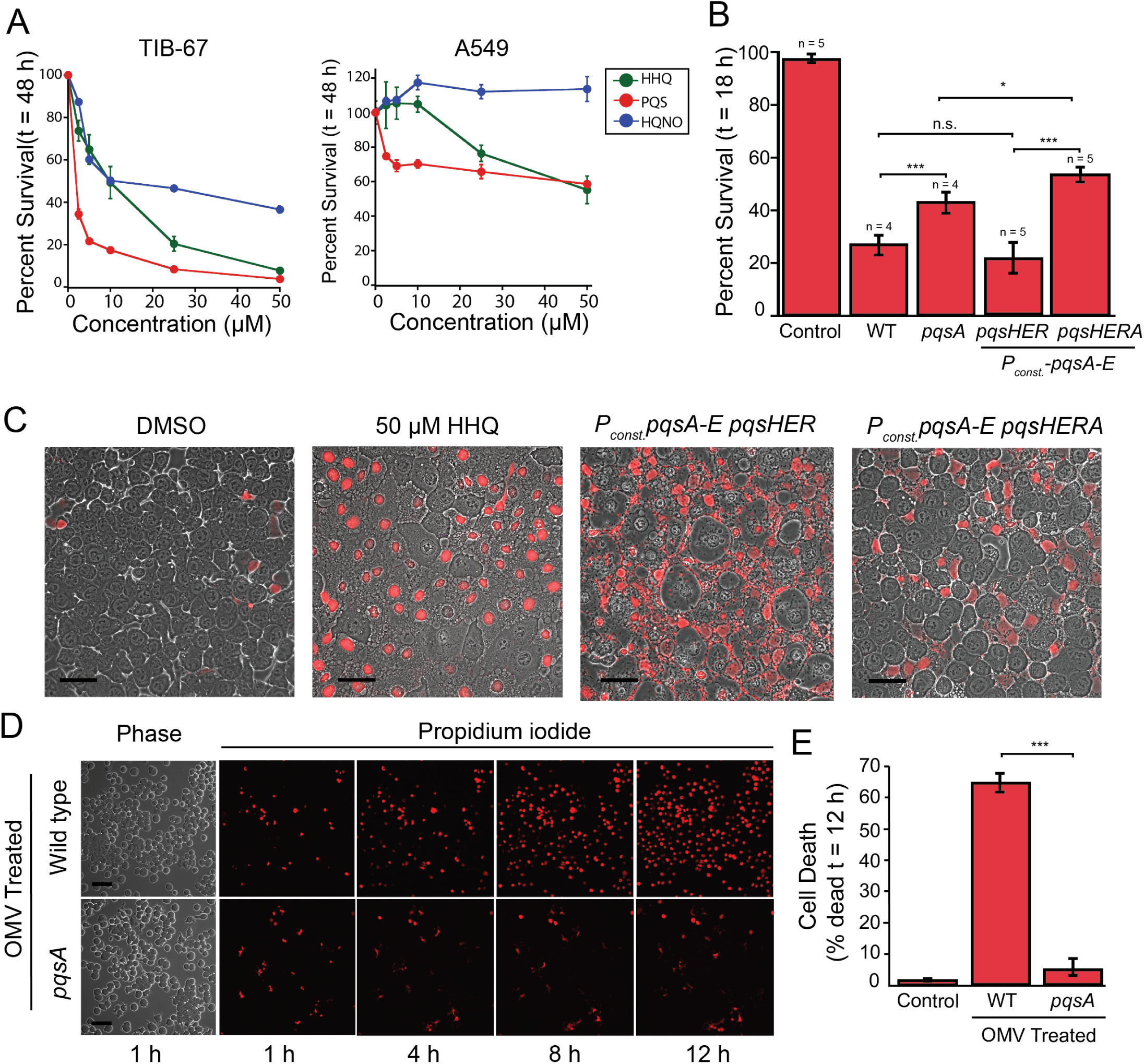
Cytotoxicity of alkyl-quinolones to TIB-67 mouse monocytes and human A549 epithelial cells. (A) MTT cell viability assay of A549 human bronchial epithelial cells or TIB-67 mouse monocytes after 48 h of treatment with various concentrations of the alkyl-quinolones HHQ, PQS, or HQNO in a 96-well format. Percent survival is relative to an untreated control. Values are mean ± SEM of three biological replicates (n = 3). (B) Percent survival of TIB-67 monocytes after 18 h co-cultured with surface-attached *P. aeruginosa*. Values are mean ± SEM of five biological replicates (n = 5). (C) Representative images of TIB-67 monocytes co-cultured with surface-attached *P. aeruginosa* or treated with exogenous HHQ under conditions of the microscopy-based virulence assay. Propidium iodide (PI) staining of DNA of non-viable cells indicates cell death (scale bars = 50 *μ*m). (D) Representative images of TIB-67 monocytes following treatment with outer membrane vesicles (OMV) purified from wild type or Δ*pqsA P. aeruginosa* culture supernatants. (E) Quantification of OMV toxicity towards TIB-67 monocytes. Percent death is the increase in PI-stained nuclei divided by the total PI-negative cells at 1 h. Values are mean ± SEM of three biological replicates (n = 3). Approximately 2500-3000 cells were analyzed for each measurement in (B) and (E). Statistical analysis in (B) and (E) is Student’s t-test (two-tailed, * = p<0.05, ** p <0.01, ***p<0.001) **Figure 4–Figure supplement 1.** Schematic of the monocyte cell death assay.

We also developed an imaging-based virulence assay adapted from previous studies (***Siryaporn et al., 2014***) (Figure 4-S1) to determine if AQs are necessary for surface-induced *P. aeruginosa* virulence towards mammalian cells. Comparing the wild type and Δ*pqsA P. aeruginosa* revealed significantly more surface-induced monocyte killing by the wild type bacteria (Figure 4B). To more precisely distinguish the effect of AQ cytotoxicity from other activities mediated by PQS, we assayed the virulence of *pqsH pqsE pqsR P*_*const*_ − *pqsABCDE* and an isogenic, *pqsA* null, derivative (Figures 4B and 4C). In the absence of PqsH, PqsE, and PqsR, any difference in virulence between these strains should be solely dependent on the ability of the bacteria products of PqsABCD such as HHQ. The strain producing HHQ caused significantly more killing than the *pqsA* mutant (Figures 4B), indicating that PQS-independent AQ activity increases the cytotoxicity of *P. aeruginosa* towards TIB-67 mouse monocytes. Thus, AQs represent important cytotoxins that promotes the surface-induced ability of *P. aeruginosa* to kill both amoebae and monocytes and this activity does not require PQS. The fact that loss of AQ production completely eliminates surface-dependent virulence towards amoebae (Figure 1C) but not towards monocytes (Figure 4B) indicates that there are also additional factors that can partially compensate for the loss of AQs in the context of monocyte infection.

An intriguing and poorly understood feature of *P. aeruginosa* virulence is the ability of these bacteria to secrete cytotoxic outer membrane vesicles (OMVs), packed with multiple virulence factors (***Bomberger et al., 2009***). PQS stimulates OMV production and is known to be abundant in OMVs (***Mashburn-Warren et al., 2009***). Given our identification of AQs as potent cytotoxins, we hypothesized that they could play a role in mediating both OMV production and cytotoxicity. We therefore treated monocytes with equal amounts of OMVs purified from wild type or Δ*pqsA P. aerguinosa*, and monitored monocyte death (Figure 4D). OMVs containing AQs were significantly more cytotoxic than those that did not (Figure 4E), indicating that AQs are a major contributor to the cytotoxicity of OMVs, not just their production.

## Discussion

Multiple lines of evidence support our conclusion that host-cell killing in response to *P. aeruginosa* surface association is mediated by induction of AQ cytotoxins. Loss of PqsABCDE, which inhibits AQ production, leads to loss of surface-induced virulence towards *D. discoideum* and significantly reduces the surface-induced virulence towards monocytes. Thus, AQ production is important for surface-induced virulence towards multiple host types. Restoring PqsABCD in the absence of PqsE, PqsH, or PqsR rescues surface-induced virulence, indicating that a PQS-independent AQ, such as HHQ, is sufficient to induce virulence even in the absence of genes responsible for AQ-mediated QS or iron-dependent signaling. Finally, purified HHQ and PQS are sufficient to directly kill host cells in the absence of bacteria, indicating that these factors are themselves cytotoxins as opposed to regulators of additional factors. These studies reinforce recent suggestions that AQs serve multiple important functions in virulence (***Lin et al., 2018***) and add surface-induced host cytotoxins to this growing list.

Our findings also implicate AQ production as a powerful reporter for dissecting the signal transduction pathways responsible for surface-induced virulence. Our analysis of the two known regulators of surface-induced virulence in the *Dictyostelium* cytotoxicity assay, PilY1 and LasR, showed that both regulators control virulence by activating *pqsABCDE* expression. Specifically, deletion of *pilY1* and *lasR* reduced *pqsA* promoter activity, PQS production, and surface-induced virulence. Meanwhile, constitutive expression of *pqsABCDE* was sufficient to restore the surface virulence of *pilY1* and *lasR* mutants and to increase the virulence of planktonic cells. Epistasis studies on both virulence and AQ production revealed that the repression of AlgR by PilY1 promotes high levels of *pqsABCDE* expression in surface-attached cells. A similar epistatic relationship between PilY1 and the AlgR-AlgZ system was recently implicated in the virulence of *P. aeruginosa* towards *C. elegans* hosts (***Marko et al., 2018***), suggesting that this pathway is also important in other virulence contexts. In the future it will be important to determine how AlgR/Z directly or indirectly regulate *pqsABCDE* expression. Additional outstanding questions include how LasR- and PilY1-dependent signaling is actually modulated by surface association and how these pathways intersect with other surface-dependent signaling pathways such as those mediated by TFP/cAMP or c-di-GMP (***Bohn et al., 2009***; ***Luo et al., 2015***; ***Persat et al., 2015***; ***Laventie et al., 2019***).

More than 50 distinct AQs have been identified in *P. aeruginosa* cultures (***Lépine et al., 2004***). Historically, the bulk of the work on AQs has focused on PQS and its roles in quorum sensing and the iron starvation response (***Lin et al., 2018***). More recently, a greater appreciation has emerged that AQs can carry out diverse functions that range from signaling (***Lee and Zhang, 2015***; ***Rampioni et al., 2016***), iron scavenging (***Diggle et al., 2007***), antibacterial tolerance (***Hazan et al., 2016***), OMV induction (***Mashburn-Warren et al., 2009***), immune suppression (***Kim et al., 2010***), and cytotoxicity towards host cells or other bacteria (***Abdalla et al., 2017***; ***Wu and Seyedsayamdost, 2017***). Our work supports this expanded view of the functional repertoire of AQs, representing evidence that AQ production is important for surface-induced virulence and demonstrating that HHQ can function as a cytotoxin that directly kills host cells independently of PQS.

The fact that AQs are sufficient to explain surface-induced *D. discoideum* killing was surprising because *P. aeruginosa* is generally thought to kill hosts using a large and redundant set of cytotoxins (***Valentini et al., 2018***; ***Streeter and Katouli, 2016***). The large number of toxins present in *P. aeruginosa* could be due to the preferential ability of different toxins to kill different host cell types. For example, A549 epithelial cells appeared less sensitive to AQ cytotoxicity in our assays, such that other cytotoxins like T3SS could take precedence in targeting these cells. While future studies will be needed to dissect the specific contributions of each virulence factor in different contexts such as with different hosts and in the presence of an intact immune system, our study highlights the value of quantitative assays that can define the specific capacities of different factors.

*D. discoideum* cells are highly phagocytic, suggesting that host delivery might be a limiting step given the poor solubility of AQs. Our data suggest that HHQ and PQS are more active when delivered to hosts by bacteria, as the AQ biosensor indicated that lower levels of HHQ are required to kill *D. discoideum* when the HHQ is produced by bacteria than when supplemented exogenously. The increased cytotoxicity of AQs when produced by bacteria could be related to OMVs. PQS is a strong stimulator of OMV production (***Mashburn-Warren et al., 2009***), and our work shows that AQs are necessary for OMV toxicity. OMVs could thus represent a built-in cytotoxin delivery system that increases the ability of *P. aerguniosa* to use AQs to both promote virulence and mediate intercellular signaling.

Given that multiple AQs can act as cytotoxins, which AQ species is likely to mediate host cell killing in vivo? HHQ and other AQs are present in the serum, urine, and sputum of CF patients with *P. aeruginosa* infections (***Collier et al., 2002***; ***Barr et al., 2015***), and have been shown to correlate with clinical progression (***Barr et al., 2015***). The conversion of HHQ to downstream AQs requires oxygen (***Schertzer et al., 2010***; ***Recinos et al., 2012***), and many infection sites are microaerophilic (***Kamath et al., 2017***; ***Hassett et al., 2009***). Indeed, HHQ levels were found to be considerably higher than PQS or HQNO in a mouse burn infection model (***Xiao et al., 2006a***). Since bacterial biofilms on surfaces such as the CF lung are also oxygen-limited (***Kamath et al., 2017***; ***Hassett et al., 2009***), HHQ may warrant particular attention as a candidate AQ cytotoxin in these conditions. In addition to being oxygen-insensitive, HHQ is a poor agonist of PqsR, with 100-fold weaker activation than PQS (***Xiao et al., 2006a***) (also see Figure 3A). Consequently, blocking the conversion of HHQ to PQS could represent a mechanism for *P. aeruginosa* to specifically induce a cytotoxic AQ without inducing competing PQS-dependent targets such as pyocyanin. Given that PQS is the most potent PqsR agonist among AQs (***Xiao et al., 2006a***) and that a recent study found that HQNO is the most potent antibiotic (***Thierbach et al., 2017***), we suggest that AQs could be functionally specialized with PQS serving primarily as a QS signaling molecule, HQNO primarily for inter-bacterial competition, and HHQ for host cell cytotoxicity under oxygen-limited conditions. Finally, we note that the different activities of AQs are not mutually exclusive such that induction of AQs upon surface association could represent a powerful strategy to simultaneously initiate cytotoxicity to ward off engulfment by phagocytic cells, suppress immune function (***Kim et al., 2010***), and signal additional downstream factors to promote factors associated with later stages of infection (***Rampioni et al., 2016***).

## Methods and Materials

### Bacterial Strains, Plasmids, and Growth Conditions

The strains and plasmids used in this study are described in Tables 1 and 2. Bacterial cultures were routinely grown in lysogeny broth (LB) broth at 37°C with aeration or on LB solidified with 1.5% agar (BD Biosciences, San Jose, CA). When stated, bacteria were grown in PS:DB media, which consists of development buffer (DB) (5 mM KH_2_PO_4_, 2 mM MgCl_2_, pH 6.5) and 10% (v/v) PS medium (10 g L^−1^ Special Peptone (Oxoid, Hampshire, United Kingdom), 7 g L^−1^ Yeast Extract (Oxoid, Hampshire, United Kindom), 10 mM KH_2_PO_4_, 0.45 M Na_2_HPO_4_, 15 g L^−1^ glucose, 20 nM vitamin B12, 180 nM Folic Acid, pH 6.5). Antibiotics were added at the following concentrations when appropriate: carbenicillin 300 μg mL^−1^, gentamycin 30 μg mL^−1^, and tetracycline 200 μg mL^−1^ for *P. aeruginosa*; 100 μg mL^−1^, gentamycin 30 μg mL^−1^, and tetracycline 15 μg mL^−1^ for *E. coli*. Expression of P_*tac*_ - or P_*lac*_ -controlled genes was induced with 1 mM IPTG. When indicated, cultures were supplemented with HHQ, PQS, or HQNO (Cayman Chemicals, Ann Arbor, MI). Unless otherwise stated, chemicals and reagents were purchased from Sigma Aldrich (St. Louis, MO).

**Table 1.** Bacterial strains and cell lines used in this study.

| Parent Strain | Strain | Relevant Characteristics | Source |
| --- | --- | --- | --- |
| *P. aeruginosa* PA14 | Wild type | - | *Rahme et al. (1995)* |
| | *lasR* | $\Delta lasR$ | *OLoughlin et al. (2013)* |
| | *pilY1* | $\Delta pilY1$ | *Kuchma et al. (2015)* |
| | *pqsA* | $\Delta pqsA$ | This study |
| | *pqsE* | $\Delta pqsE$ | This study |
| | *pqsH* | $\Delta pqsH$ | This study |
| | *pqsR* | $\Delta pqsR$ | This study |
|  | *pqsL* | *pqsL::Mar2xT7* | *Liberati et al. (2006)* |
| | *algR* | $\Delta algR$ | This study |
| | *algR pilY1* | $\Delta algR \Delta pilY1$ | This study |
| | *phz* | $\Delta phzABCDEF G1 \Delta phzABCDEF G2$ | This study |
|  | *rhlB* | *rhlB::Mar2xT7* | *Liberati et al. (2006)* |
|  | *hcnB* | *hcnB::Mar2xT7* | *Liberati et al. (2006)* |
| | *rsmA* | $\Delta rsmA$ | This study |
| | *gacA* | $\Delta gacA$ | This study |
| | *vfr sadC* | $\Delta vfr \Delta sadC$ | *Merritt et al. (2010)* and this study |
| | *pvdS fpvI* | $\Delta pvdS \Delta fpvI$ | This study |
| | $P_{const} - pqsA-E$ | $P_{OXB20}$ | This study |
| | $P_{const} - pqsA-E pqsR$ | $P_{OXB20}::pqsA \Delta pqsR$ | This study |
| | $P_{const} - pqsA-E pqsH$ | $P_{OXB20}::pqsA \Delta pqsH$ | This study |
| | $P_{const} - pqsA-E pqsHER$ | $P_{OXB20}::pqsA \Delta pqsH \Delta pqsE \Delta pqsR$ | This study |
| | $P_{const} - pqsA-E pqsHERA$ | $P_{OXB20}::pqsA \Delta pqsH \Delta pqsE \Delta pqsR pqsA::Mar2xT7$ | This study |
| | $P_{const} - pqsA-E lasR$ | $P_{OXB20}::pqsA \Delta lasR$ | This study |
| | $P_{const} - pqsA-E pilY1$ | $P_{OXB20}::pqsA \Delta pilY1$ | This study |
| | $P_{pqsA} - mCherry$ | *attB::P<sub>pqsA</sub> - mCherry* | This study |
| | $P_{pqsA} - mCherry pilY1$ | *attB::P<sub>pqsA</sub> - mCherry $\Delta pilY1$* | This study |
| | $P_{pqsA} - mCherry lasR$ | *attB::P<sub>pqsA</sub> - mCherry $\Delta lasR$* | This study |
| | $P_{lac} - vector$ | pBBRMCS3 | *Kovach et al. (1995)* and this study |
| | $P_{lac} - algRD54E$ | pBBRMCS3::*P<sub>lac</sub> - algRD54E* | This study |
| *P. aeruginosa* PAO1 | AQ Biosensor | $\Delta pqsA \Delta pqsR glmS::P_{tac} - pqsR$<br>*pUCP18::P<sub>rpoD</sub> - mKate P<sub>pqsA</sub> - YFP* | This study |
| *E. coli* B/r | Wild type | - | *Siryaporn et al. (2015)* |
| *D. discoideum* | AX3 | - | *Loomis (1971)* |
| *M. musculus* | J774A.1(ATCC® TIB-67™) | - | ATCC |
| *H. sapiens* | A549 (ATCC® CCL-185™) | - | ATCC |

**Table 2.** Plasmids used in this study.

| Parent Strain | Plasmid | Source |
| --- | --- | --- |
| E. coli S17 | pEXG2 | *Hmelo et al. (2015)* |
|  | mini-CTX-2 | *Hoang et al. (2000)* |
|  | pUC18-mini-Tn7T-Gm | *Choi and Schweizer (2006)* |
|  | pUC18-mini-Tn7T-LAC | *Choi and Schweizer (2006)* |
|  | pUCP18-RedS | *Lesic and Rahme (2008)* |
|  | pFLP2 | *Hoang et al. (1998)* |
| | pUCP18::$P_{rpoD}$ – mKate $P_{Paqa}$ – YFP | *Persat et al. (2015)* |
| | mini-CTX-2::$P_{tac}$ – mCherry | *Siryaporn et al. (2015)* |
|  | pBBRMCS3 | *Kovach et al. (1995)* |
|  | pSF-OXB20 | Oxford Genetics |
|  | pUC19 | New England Biolabs |

### Strain Construction

Primers used in this study are described in Table 3. All gene deletions described here are unmarked, in-frame deletions generated by two-step allelic exchange, as described previously (***Hmelo et al., 2015***). Briefly, upstream and downstream homology arms flanking the relevant gene were amplified with primer pairs (-KO1,-KO2 and -KO3,-KO4; Table 3), fused through overlap extension PCR (OE-PCR), and cloned into restriction sites of plasmid pEXG2. The pEXG2 plasmid was integrated into *P. aeruginosa* through conjugation using the donor strain *E. coli* S17. Exconjugants were selected on gentamycin and then the mutants of interest were counterselected on 5% sucrose. Transposon insertions obtained from the PA14 Transposon Mutant Database (***Liberati et al., 2006***) were transferred between strains using the *λ*-Red recombination system (***Lesic and Rahme, 2008***).

**Table 3.**
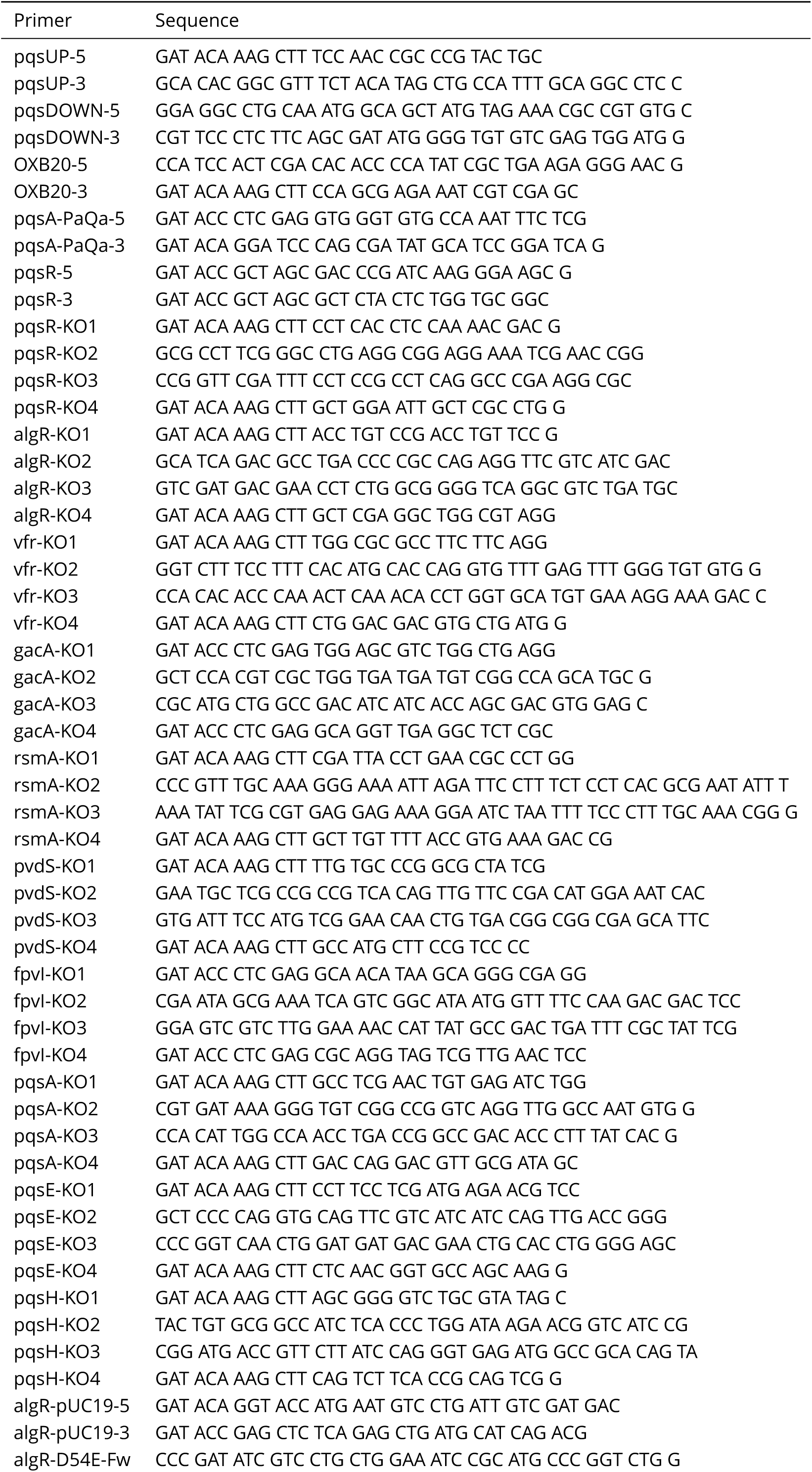
Primers used in this study.

To generate the *pqsABCDE* overexpression strain, the *P*_*OXB20*_ promoter was amplified from the plasmid pSF-OXB20 (Oxford Genetics, Cambridge, MA) using primer pair (OXB20-5 and OXB20-3) and spliced between two 400 bp fragments that flank the *pqsA* promoter, which were amplified from gDNA using primers (pqsUP-5, pqsUp-3) and (pqsDOWN-5, pqsDOWN-3), respectively. The resulting construct was cloned into pEXG2 and integrated onto the chromosome using allelic exchange. To generate the inducible *pqsR* expression construct, the *pqsR* gene was amplified from gDNA with primer pair (pqsR-5, pqsR-3) and cloned into pUC18-mini-Tn7T-LAC. Proper gene orientation was confirmed by restriction mapping, and the resulting construct was integrated onto the chromosome by co-electroporation with pTNS2 using methods described previously (***Choi and Schweizer, 2006***).

To generate the *algRD54E* mutation and overexpression construct, the *algR* gene was amplified from gDNA using primers (algR-pUC-5, algR-pUC-3) and subcloned into pUC19 (New England Biolabs, Ipswich, MA). Site-directed mutagenesis was performed by amplification of pUC19::*algR* with primers (algR-D54E-Fw, algR-D54E-Rv), and the mutant allele was either cloned into pEXG2 and integrated into the *P. aeruginosa* chromosome by allelic exchange, or cloned into pBBRMC3 to obtain the overexpression construct. The *P*_*pqsA*_ − *mCherry* promoter fusion construct was generated by amplifying an approximately 500 bp fragment upstream of the *pqsA* gene with primer pair (P1pqsA-1, P1pqsA-2) and fusing it by OE-PCR to a promoterless *mcherry* gene, amplified from mini-CTX-2::PA1/04/03-mCherry with primer pair (P1pqsA-3, P1pqsA-4). The resulting fragment was cloned into the mini-CTX-2 plasmid and integrated into the chromosome at the CTX-2 phage attachment site (attB).

To generate the fluorescent AQ biosensor, a fragment containing the *pqsA* promoter and PqsR binding site (−19 bp to −219 bp) was amplified from gDNA using primer pair (pqsA-PaQa-5, pqsA-PaQa-3), and cloned into the BamHI/XhoI sites of pUCP18::*P*_*pqsA*_ − *YFP P*_*rpoD*_ − *mKate*. This plasmid was then transformed into a PAO1 strain containing a deletions in *pqsA*, and replacement of the *pqsR* promoter with a constitutive *P*_*tac*_ promoter. We note that while our virulence assays were all performed with the PA14 strain of *P. aeruginosa*, we used the PAO1 strain for the biosensor since the reporter is avirulent and does not interfere with PA14 virulence while PAO1 maintains plasmids at higher levels than PA14.

### *D. discoideum* and mammalian cell culture

*D. discoideum* AX3 was maintained axenically as described previously (***Siryaporn et al., 2014***; ***Fey et al., 2007***). Briefly, frozen stocks were inoculated into overnight cultures of *E. coli* B/r, and plated on GYP plates. After incubation for 4-6 days at 22°C, individual spores were inoculated into PS media supplemented with Antibiotic-Antimycotic (AA) solution, and incubated at 22°C 100 rpm. When cultures reached approximately 1 × 106 cells mL^−1^, cells were back-diluted 1:100 in fresh PS media. Axenic cultures were maintained for up to 1 month.

J774A.1 mouse monocytes (ATCC® TIB-67) and A549 (ATCC® CCL-18) (verified mycoplasma free) were grown at 37°C with 5% CO_2_ in Dulbecco’s Modified Eagle’s Medium (Gibco, Dublin, Ireland) supplemented with 10 % fetal bovine serum and Penicillin-Streptomycin solution (Invitrogen, Grand Island, NY). Cells were passaged according to the ATCC protocols.

### *D. discoideum* cell death assay

Cell death assays were performed as described previously with minor modifications (***Siryaporn et al., 2014***). Overnight cultures of *P. aeruginosa* were diluted 1:100 in PS:DB media and grown to OD_600*nm*_ = 0.6-0.8 at 37 °C with aeration. Cultures were transferred to glass-bottom dishes (Mattek, Ashland, MA) and incubated for an additional 1 hour at 100 rpm on a rotary shaker. For the planktonic condition, aliquots of culture media were washed with PS:DB, concentrated 20-fold, and plated onto fresh glass-bottom dishes. For the surface-attached condition, culture media was aspirated and surface-attached cells were washed with PS:DB. Aliquots of *D. discoideum* culture (between 2-5 × 10^5^ cells mL^−1^) were washed with PS:DB, and added to planktonic and surface-attached bacteria to achieve the appropriate multiplicity of infection (MOI), which ranged between roughly 500:1 to 1000:1 (*P. aeruginosa* to *D. discoideum*), unless otherwise stated. See ***Siryaporn et al.*** (***2014***) for details regarding assay validation and MOI quantification. The combined samples were covered with a 1% agar pad, prepared by pouring molten 1% agar in PS:DB containing 1 *μ*M calcein-AM (Invitrogen, Grand Island, NY) on a glass surface, and cutting the solidified pad into 1 cm x 1 cm sections. Samples were analyzed by imaging cells with phase contrast and FITC channels after 1 hour of incubation at 25 °C using fluorescence microscopy. Cell death was quantified by counting the total number of calcein-AM-positive and -negative cells. All reported values of percent cell death are averages of at least three independent experiments (biological replicates). Each set of experiments included wild type and *pqsA* mutant controls, and each measurement was of 150-500 cells.

For quantifying cytotoxicity of purified AQs under conditions of the microscopy-based cell death assay (Figure 1D), AQs were diluted 1:200 into molten 1% agar in PS:DB and pads were prepared as described above. Samples of *D. discodium* were transferred to glass-bottom dishes and covered with a 1 cm × 1 cm agar pad and incubated at 25°C for 1 hour. For quantifying cytotoxicity of purified AQs in axenic *D. discoideum* cultures (Figure 1E), *D. discoideum* was subcultured to 10,000 cells mL^−1^ in fresh PS media with antibiotics and varying concentrations of AQs. Cultures were incubated at 22°C with shaking at 450 rpm, and cell density was measured by counting with a hemocytometer. Experiments were performed with five biological replicates.

### Quantification of fluorescent reporter expression

Reporter strains were grown according to the procedures of the *D. discoideum* cell death assay. Bacterial cultures were transferred to glass-bottom dishes when OD_600*nm*_ reached 0.6, and incubated at 37°C with shaking (100 rpm) for 1 h. Cells were washed and isolated as described above, and single cells were imaged immediately after addition of the 1% agar pad using fluorescence microscopy. Mean fluorescence intensity per cell was computed for 500-1000 cells, and average values were normalized by the expression of a constitutive *P*_*tac*_ − *mCherry* reporter, which was measured in the same manner as the *P*_*pqsA*_) reporters. All reported values are averages from at least three independent experiments (biological replicates).

### LC/MS-based quantification of AQ production

Overnight cultures of *P. aeruginosa* were subcultured 1:100 into 20 mL PS:DB and grown to OD_600*nm*_ = 1.5 at 37°C. Cultures were extracted with equal volumes of ethyl acetate, dried and resuspended in methanol. For validations of AQ levels in *P*_*constitutive*_ − *pqsABCDE* stains shown in Figure 1-S5, cultures were grown for 18 hr to late stationary phase in LB. Samples were analyzed by HPLC-MS on a 1260 Infinity Series HPLC system (Agilent, Santa Clara, CA) equipped with an automated liquid sampler, a diode array detector, and a 6120 Series ESI mass spectrometer using an analytical Luna C18 column (5 m, 4.6 x 100 mm, Phenomenex, Torrance, CA) operating at 0.6 mL min^−1^ with a gradient of 25% MeCN in H_2_O to 100% MeCN over 18 minutes. A standard curve constructed from commercial AQ standards was used to calculated AQ concentrations in cultures.

### Plate-based AQ biosensor assay

Overnight cultures of the AQ biosensor was diluted 1:100 in LB with 1 mM IPTG and grown to late exponential phase (OD_600*nm*_ = 1.0-2.0). Bacteria were resuspended in fresh LB to OD_600*nm*_ = 1.0, and added to equal volumes of LB supplemented with various concentrations of AQs (purchased from Sigma-Aldrich) in a 96-well plate. Plates were sealed with a breathable membrane (DivBio, Dedham, MA) and incubated at 37°C 450 rpm while monitoring fluorescence in the YFP (500 nm excitation/540 nm emission) and mKate (590 nm excitation/645 nm emission) channels. Biosensor signal was calculated by normalizing the YFP signal by the mKate signal and subtracting the baseline expression from a DMSO control. Six technical replicates were performed for all conditions shown in Figure 3A.

### Biosensor-based AQ quantification of surface-attached bacteria

Overnight cultures of *pqsH* mutant *P. aeruginosa* were grown following the procedures of the *D. discoideum* cell death assay. The AQ biosensor was prepared according to the procedures of the plate-based assay, but resuspended to OD_600*nm*_ = 0.2 in PS:DB. For planktonic samples, biosensor was added to equal volumes of planktonic cell suspensions. For surface-attached samples, biosensor was added to washed, surface-attached cells. Cells were covered with a 1% agar pad in PS:DB (1 mM IPTG), and incubated at 25 °C for 1-2 hours. Single-cell YFP and mKate fluorescence was measured by microscopy, and biosensor signal was calculated by normalizing the YFP signal by the mKate signal, and subtracting by the baseline expression of biosensor doped into a lawn of surface-attached Δ*pqsA P. aeruginosa*. For clarity, biosensor signal in representative microscopy images are normalized by the mean biosensor signal of biosensor doped into a lawn of surface-attached Δ*pqsA*. All reported biosensor signal measurements are averages from at least three independent experiments, and approximately 300-500 individual cells were analyzed for each condition. An HHQ standard curve was generated and used to convert biosensor signal to HHQ concentration. Biosensor was added to surface attached Δ*pqsA P. aeruginosa* and covered with 1% agar pads made with PS:DB, supplemented with various concentrations of HHQ. Biosensor signal in response to HHQ standards was calculated as described above. A new standard curve was constructed for each independent experiment. Measurements of biosensor signal of surface-attached Δ*pqsH* that fell beyond the saturation point of the HHQ standard curve were not included in the analysis to prevent over-estimation of the concentrations of HHQ in surface-attached populations.

### Mammalian cytotoxicity assay of purified AQs

TIB-67 monocytes or A549 epithelial cells were seeded at a density of 150,000 cells or 75,000 cells cm^−2^ in a 96-well plate and incubated at 37°C 5% CO_2_ for 24 hours. Cells were treated with purified AQ compounds dissolved in DMSO such that the final concentration of DMSO was less than 0.5%. After 48 hours, media was aspirated and the WST-8 reagent EZQuant (Alstem Bio, Richmond, CA) was used to assess cell viability. Values reported are averages of three biological replicates. At least three independent experiments were performed, and the trends observed in Figure 4A was observed across all experiments.

### Microscopy-based TIB-67 cell death assay

For the microscopy-based assay, TIB-67 monocytes were seeded at a density of 75,000 cells cm^−2^ and incubated at 37°C 5% CO_2_ for 24 hours. TIB-67 monolayers were washed twice with phosphate buffered saline (PBS) (Gibco, Dublin, Ireland) and combined with *P. aeruginosa* samples. To prepared *P. aeruginosa* samples, cultures were grown following the procedures of the *D. discoideum* cell death assay. When cultures reached OD_600*nm*_ = 0.5-0.6, cultures were transferred to petri dishes coated with a thin layer of 1% agar in PS:PBS (10% (v/v) PS media in PBS with 1 mM MgSO_4_ and 0.1 mM CaCl_2_, pH = 7.2) and grown for an additional 1 hour at 37°C on a rotary shaker (100 rpm) to allow surface attachment. Surface-attached bacteria were washed twice with PS:PBS. The density of surface-attached bacteria could be adjusted based on force applied during washing steps using an automated pipette, and density was optimized to achieve a final multiplicity of infection (MOI) of 1:50 to 1:150 (P. aeruginosa to TIB-67). Propidium iodide (PI;1 μM final) and sub-MIC dose of tetracycline (5 μg mL^−1^ final) was added to prepared monocyte samples. Agar pads were excised and inverted onto prepared monolayers of TIB-67 monocytes. Concentrations of PI and tetracycline were calculated based on the volume of the agar pad. Samples were incubated at 30°C in an incubated chamber, and cells were tracked over the course of 24 hours using fluorescence microscopy. Cell death was monitored by PI staining of the DNA of non-viable cells. Bacterial co-culture with monocytes resulted in substantial monocyte lysis (see Figures 4C) and subsequent diffusion of the nuclear PI stain. Therefore, *P. aeruginosa* virulence was most precisely quantified by counting viable monocytes (PI-negative) as opposed to dead monocytes (PI-positive). PI-negative cells were counted at various timepoints and divided by the number of PI-negative cells at 1 h in the same field of view. Treatment of monocytes with AQs or purified in OMVs did not result in lysis (see Figures 4C and 4D), and therefore virulence was measured by counting dead, PI-positive cells (Figure 4E).

### OMV isolation and quantification

Outer membrane vesicles (OMV) were isolated as described previously (***Bauman and Kuehn, 2006***). Because OMVs are not significantly produced in the absence of PQS, we used the approach of ***Bauman and Kuehn*** (***2006***) to stimulate OMV production in the *pqsA* mutant, and kept conditions the same for wild type cultures to minimize differences between samples. Briefly, overnight cultures of *P. aeruginosa* were subcultured 1:200 in LB and grown at 37°C 250 rpm to O_600*nm*_D = 0.3-0.4. Polymyxin B was added to a final concentration of 4 μg mL^−1^, and cultures were grown at 37°C 250 rpm until they reached an OD_600*nm*_ between 0.8-1.0. Culture supernatants were filtered through a 0.45 μm filter (Millpore, Burlington, MA), then centrifuged at 40,000 x g for 1 hour. Pellets were resuspended in PBS, filtered through a 0.45 μm filter, and centrifuged for an additional 1 hour at 100,000 x g. Samples were resuspended in PBS and concentrated with a 10,000 MWCO centrifugal filter device (Millipore, Burlington, MA). Filtrate was used as a vehicle control. OMV concentrations were measured by addition of the lipophilic dye FM-464 (Fisher Scientific, Hampton, NH) and measuring fluorescence (506 nm excitation/750 nm emission). The OMV concentration of the wild type sample was adjusted to match the concentration of the sample. Final OMV stocks were approximately 5000-fold concentrated compared to culture supernatant, and diluted 1:200 (based on the volume of the agar pad) into each sample of TIB-67 monocytes.

## Acknowledgments

We wish to thank members of the Gitai and Shaevitz labs for helpful discussions and comments on the manuscript, the Bassler and O’Toole labs for strains and reagents, and Ben Bratton for technical assistance and data analysis. ZG is supported by an NIH Pioneer Award DP1-AI124669 and an award from the Princeton Catalysis Initiative.

---

**Figure 1–Figure supplement 1.**
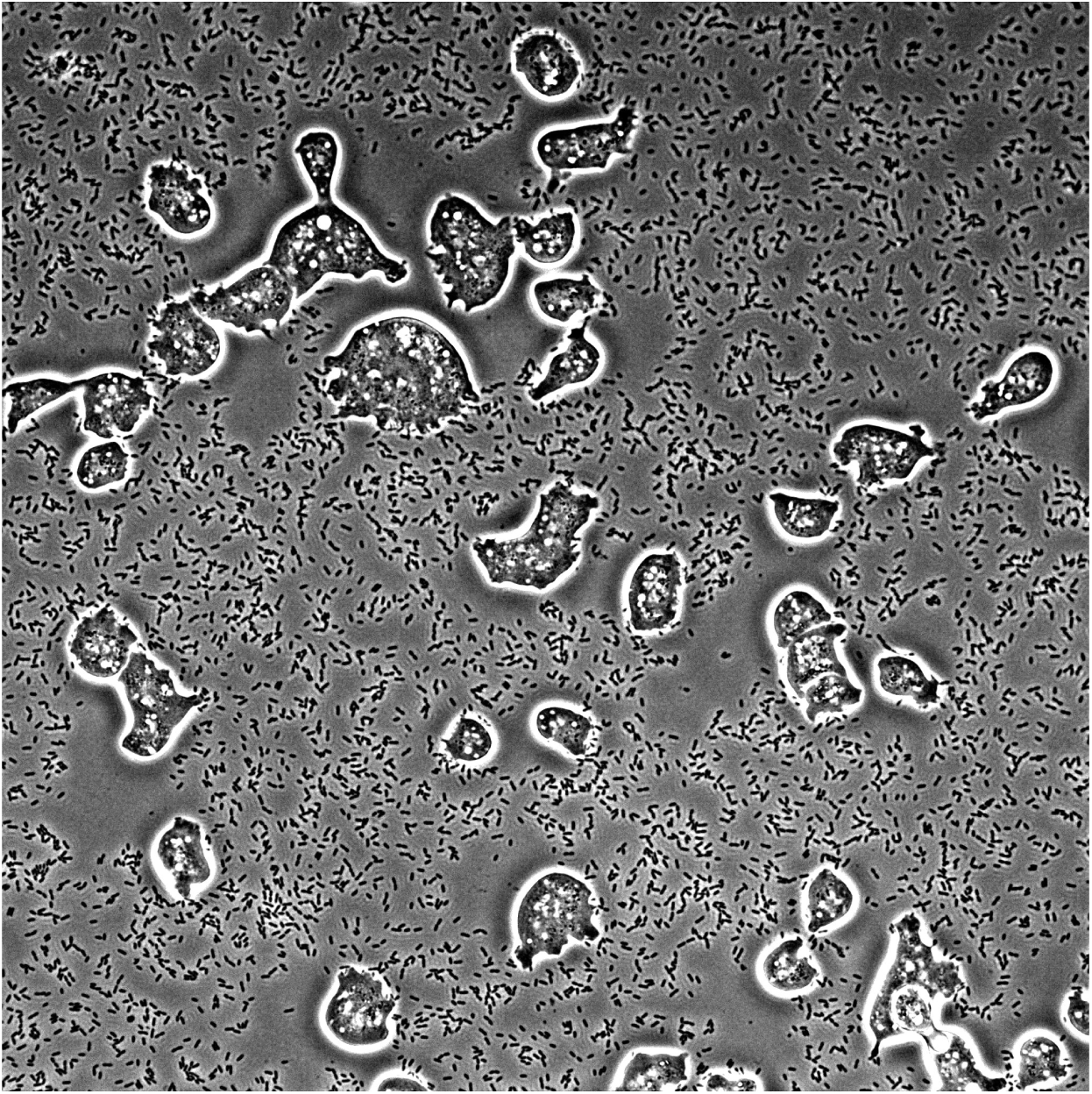
Time-lapse imaging of *D. discoideum* infected with planktonic *P. aeruginosa*. Planktonic bacteria were isolated following the procedures of the *D. discoideum* cell-death assay (see *Materials and Methods*) and added to *D. discoideum* at an MOI of approximately 50:1 (bacteria:amoeba). Samples were covered with a 1.5 % agar pad and incubated at 25°C. Phase contrast images were taken every 5 min for 200 min.

**Figure 1–Figure supplement 2.**
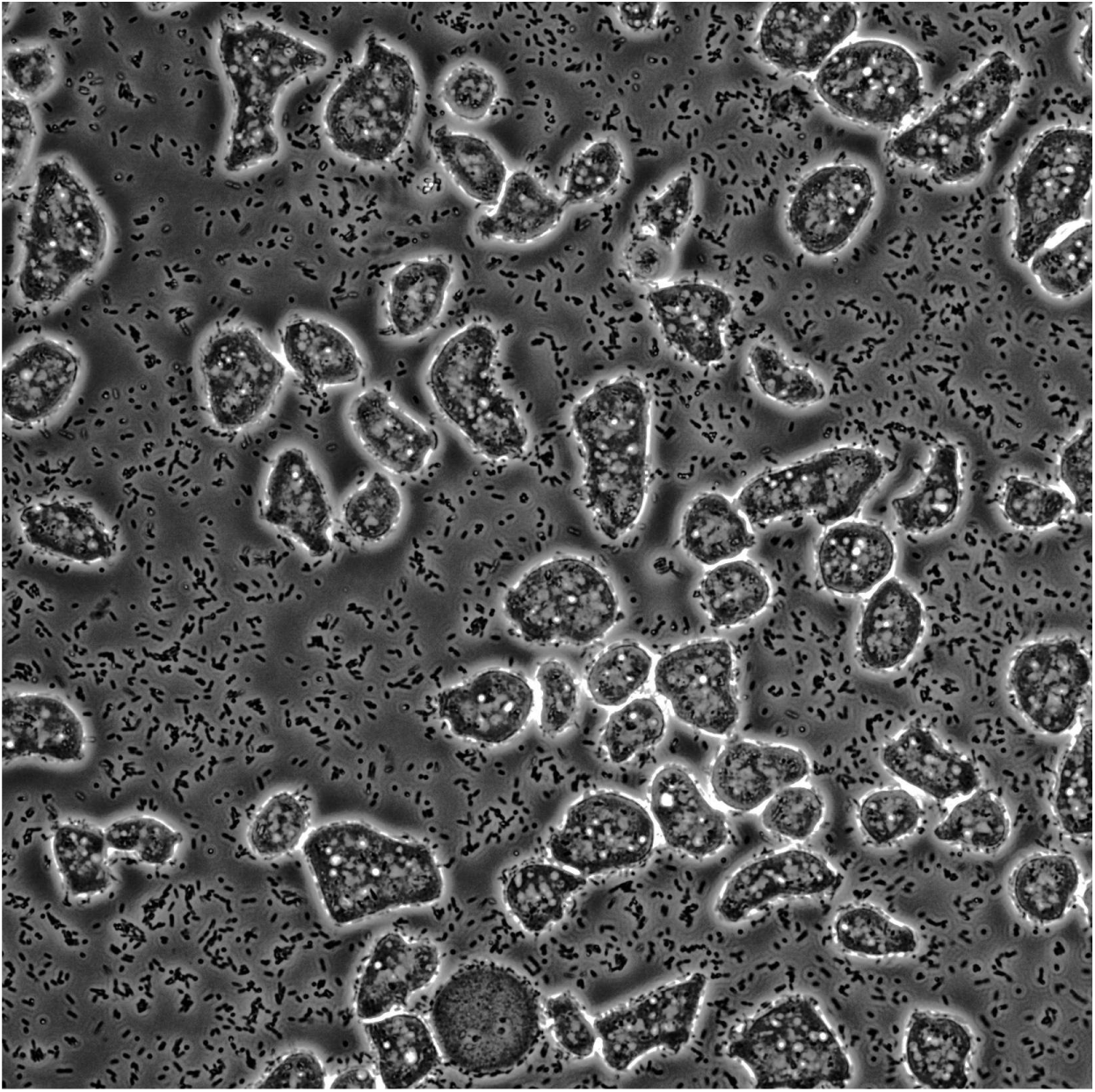
Time-lapse imaging of *D. discoideum* infected with surface-attached *P. aeruginosa*. Surface-attached bacteria were isolated following the procedures of the *D. discoideum* cell-death assay (see *Materials and Methods*) and added to *D. discoideum* at an MOI of approximately 50:1 (bacteria:amoeba). Samples were covered with a 1.5 % agar pad and incubated at 25°C. Phase contrast images were taken every 5 min for 200 min.

**Figure 1–Figure supplement 3.**
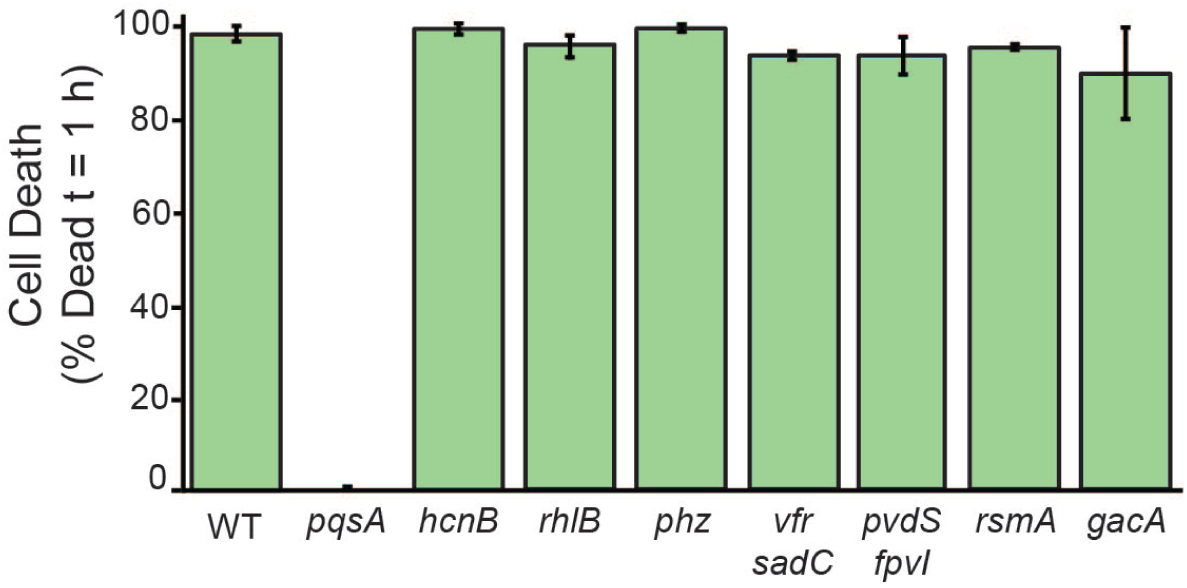
Quantification of *D. discoideum* killing by surface-attached *P. aeruginosa* mutants after 1 hour of co-culture. Cell death was indicated by positive staining by the fluorescent dye calcein-AM. Values are averages of three independent biological replicates and error bars represent standard error. Approximately 150-300 cells were analyzed for each measurement.

**Figure 1–Figure supplement 4.**
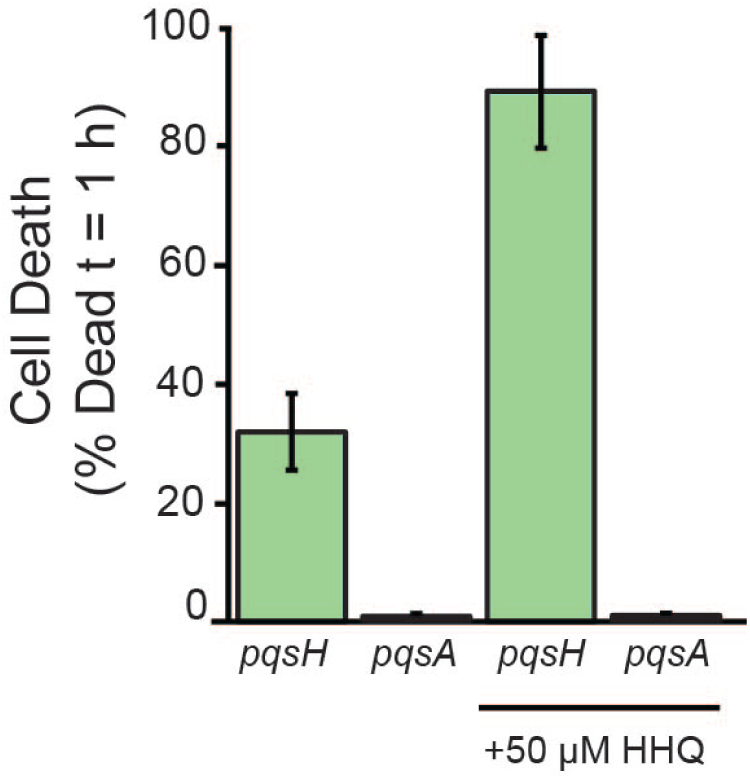
Quantification of *D. discoideum* killing by surface-attached *P. aeruginosa pqsH* and *pqsA* mutants supplemented with exogenous HHQ or PQS. HHQ, PQS, or DMSO was added to cultures after the initial 1:100 dilution, and then cultures were grown following the procedures for the D. *discoideum* cell death assay, as described in *Materials and Methods*. Surface attached and planktonic bacteria were washed twice in PSDB to remove any exogenous HHQ or PQS prior to incubating the bacteria with *D. discoideum*. Values are averages of three independent biological replicates and error bars represent standard error. Approximately 300-500 cells were analyzed for each measurement

**Figure 1–Figure supplement 5.**
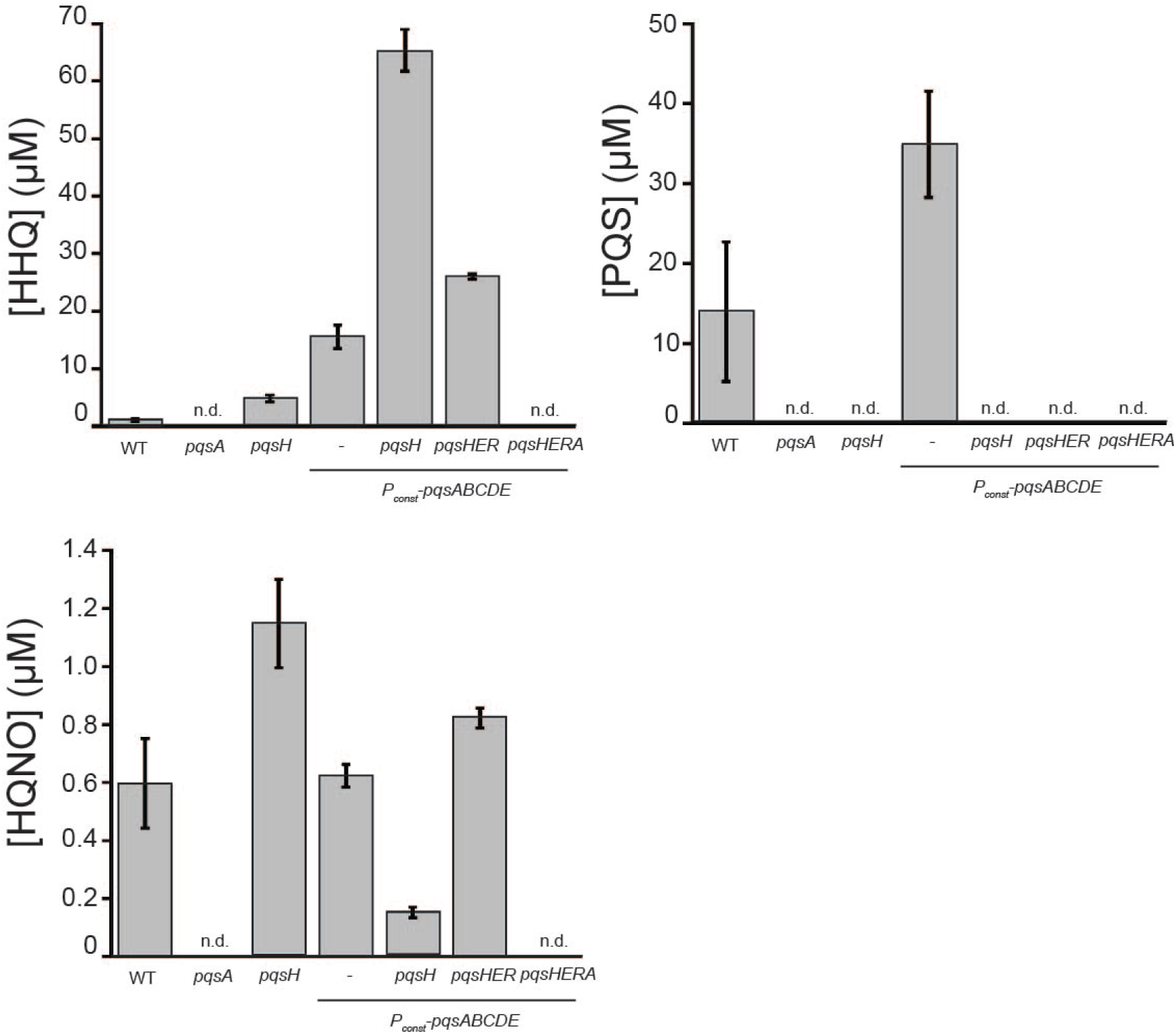
Quantification of alkyl-quinolone concentrations in wild type and mutant *P. aeruginosa* cultures using LC/MS. Stationary phase cultures were extracted with ethyl acetate, dried, and resuspended in methanol. Samples were analyzed by LC/MS, and AQ concentrations were calculated using a standard curve constructed from commercial AQ standards. Values are averages of three independent biological replicates and error bars represent standard error (n.d.=not detected).

**Figure 3–Figure supplement 1.**
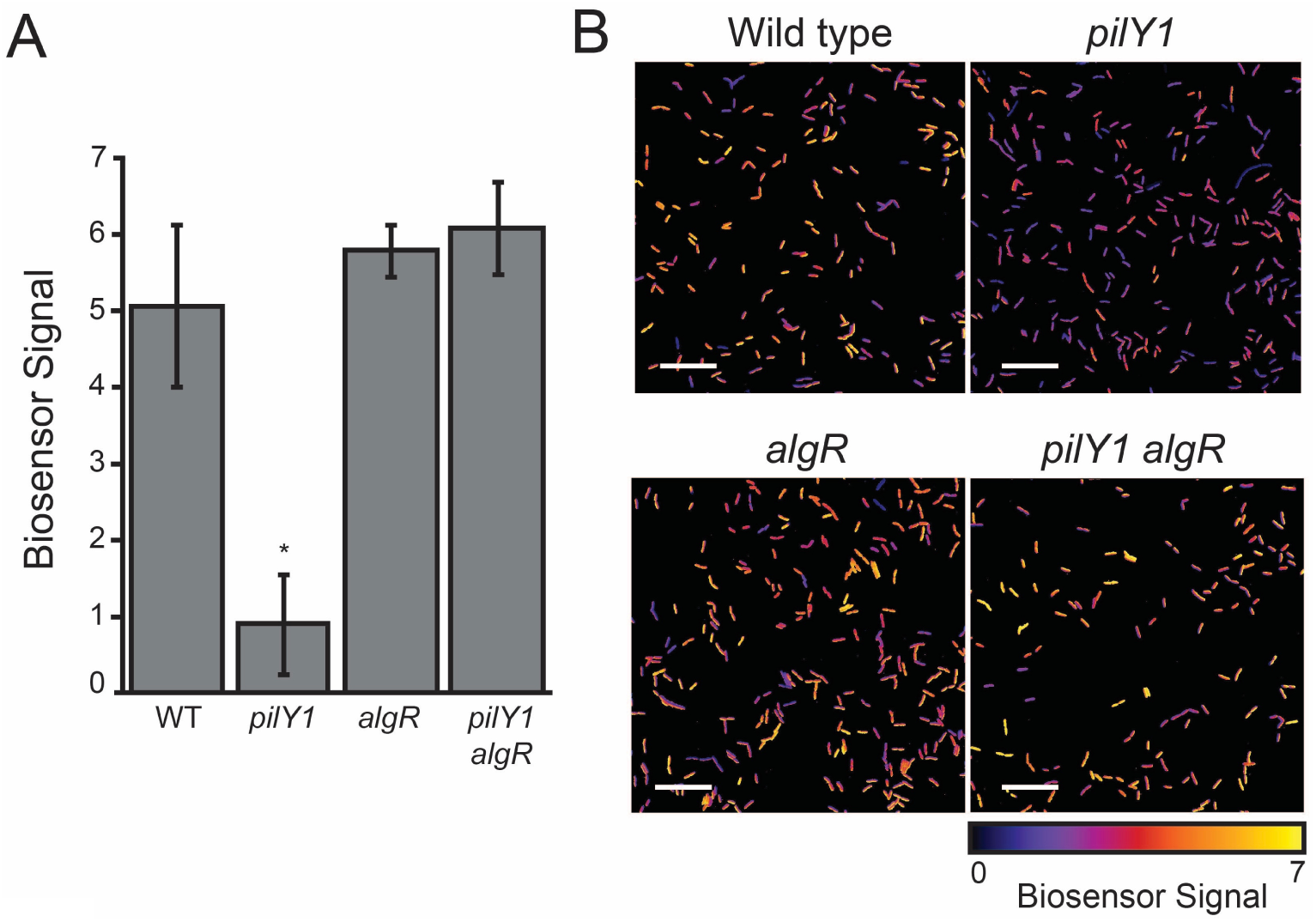
Biosensor-based quantification of AQ levels in surface-attached *P. aeruginosa* populations. AQ biosensor was doped into *P. aeruginosa* samples as described in Figure 3. Mean YFP/mKate fluorescence intensity per cell was calculated for approximately 500 cells, and values were normalized by the mean YFP/mKate fluorescence of biosensor doped into the surface-attached *pqsA* mutant. Values are averages of three independent experiments and error bars represent standard error. (B) Representative images of samples described in (A) (scale bars = 10 *μ*m).

**Figure 3–Figure supplement 2.**
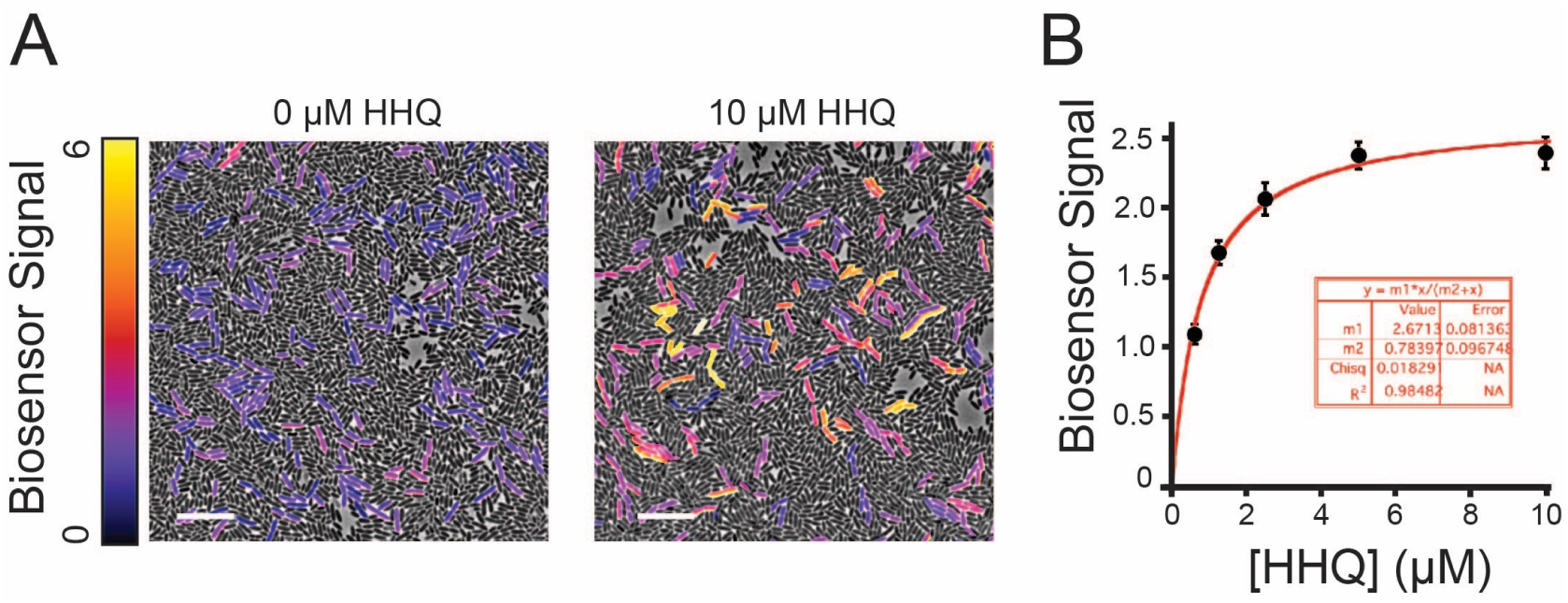
(A) Representative images of AQ biosensor doped into samples of surface-attached Δ*pqsA P. aeruginosa* treated with 10 *μ*M HHQ or DMSO added to 1% agar pads used for imaging (scale bars = 5 *μ*m). (B) Mean biosensor signal per cell calculated from approximately 500 individual cells for each HHQ concentration. Standard curves were constructed for each independent experiment and used to calculate HHQ concentrations in Figure 3. Error bars represent standard error. Curve fit was computed using KaleidaGraph software (Synergy Software, Reading, PA).

**Figure 4–Figure supplement 1.**
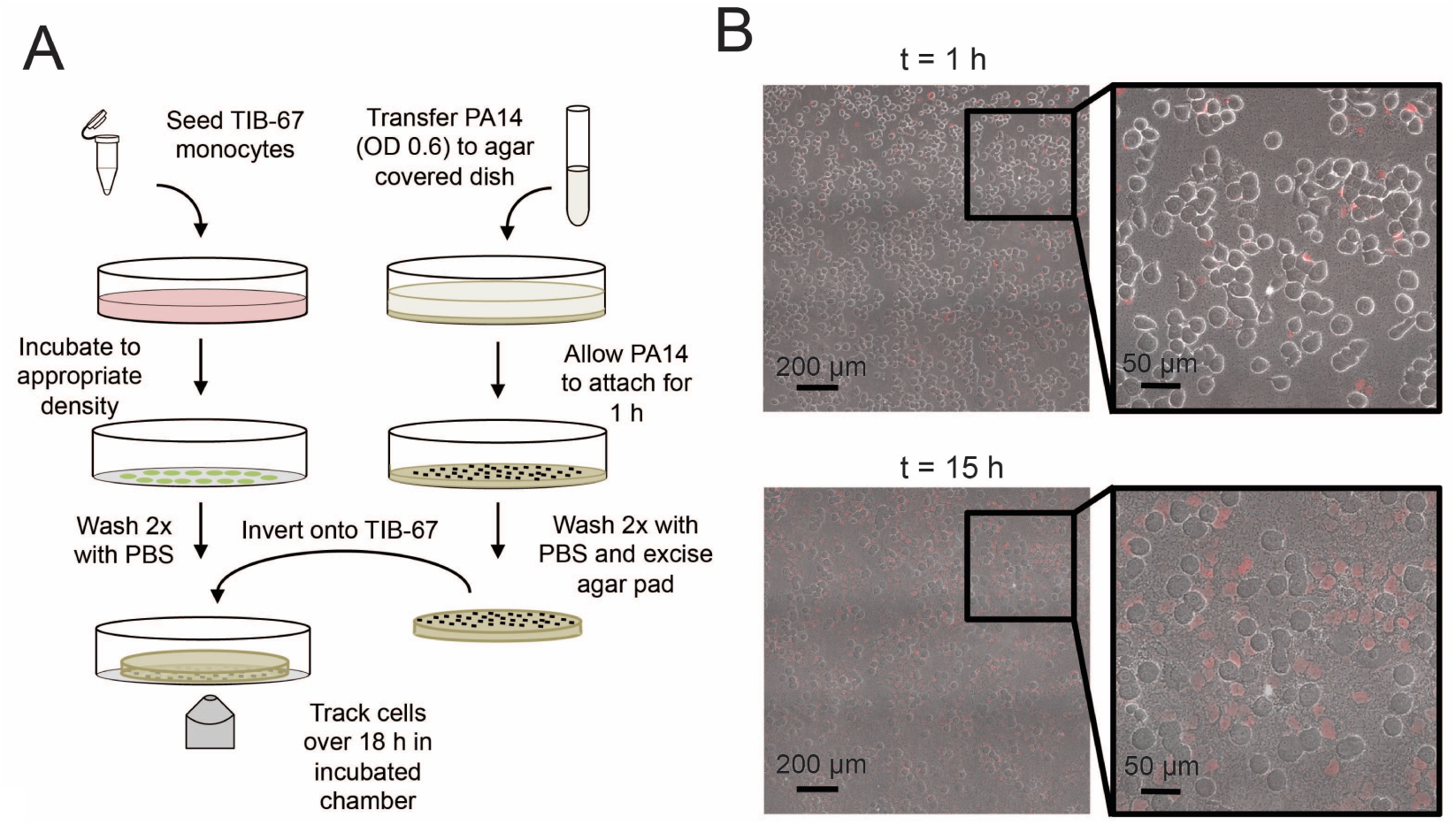
(A) Schematic of the monocyte cell death assay described in *Materials and Methods*. (B) Representative images of TIB-67 monocytes treated with surface-attached, wild type *P. aeruginosa* at a multiplicity of infection of approximately 1:50 at 1 and 15 hours of incubation at 30°C. Cell death is indicated by propidium iodide staining.

